# The regulator of G protein signaling RGS9 supports insulin secretion in male mouse islets

**DOI:** 10.64898/2026.10.26.761401

**Authors:** Laura Reininger, Sarah Ferragne, Scott A. Campbell, Caroline Tremblay, Mélanie Éthier, Julien Ghislain, Vincent Poitout

## Abstract

**Objectives:** Regulators of G protein signaling (RGS) proteins terminate G protein-coupled receptor signaling and regulate multiple aspects of pancreatic islet function. Although *RGS9* is highly enriched in human and rodent islets, its cellular localization and physiological role remain largely unknown. Here, we investigated the expression of *Rgs9* in mouse islets and its contribution to insulin secretion.

**Methods:** *Rgs9* expression was characterized by RNA in situ hybridization and fluorescence-activated cell sorting coupled with quantitative PCR. *Rgs9* function was investigated by shRNA-mediated knockdown in wild-type and Dishevelled, Egl-10 and Pleckstrin (DEP) domain-deficient *Rgs9* mutant (*Rgs9^ΔDEP/ΔDEP^*) mouse pseudo-islets.

**Results:** *Rgs9* was detected throughout the endocrine pancreas but was enriched in β and δ cells, with the *Rgs9-1* isoform representing the predominant splice variant in mouse islets. *Rgs9* knockdown in mouse pseudo-islets impaired glucose- and KCl-stimulated insulin secretion without affecting insulin content. In contrast, *Rgs9^ΔDEP/ΔDEP^* mouse islets failed to reproduce this phenotype. Further characterization of the mutant mice revealed persistent expression of *Rgs9* transcripts, which when knocked down impaired both glucose- and KCl-stimulated insulin secretion.

**Conclusions:** These data indicate that RGS9 supports glucose-stimulated insulin secretion. Given that RGS9 acts exclusively on Gαi/o, our results further suggest that RGS9 in β cells dampens inhibitory signaling to promote insulin secretion. Finally, the *Rgs9^ΔDEP/ΔDEP^* mutant does not represent a complete loss-of-function in islets likely due to the retention of the GTPase-activating protein (GAP) activity in the absence of the DEP domain.

## 1. INTRODUCTION

Islets of Langerhans express many G protein coupled receptors (GPCR) that contribute to the fine regulation of hormone secretion in response to metabolic, hormonal, and neuronal signals to maintain glucose homeostasis [1; 2]. GPCR signaling is mediated by coupling to a discrete number of heterotrimeric G proteins. Inhibitors of insulin secretion (eg. somatostatin (SST) [3], dopamine [4; 5] and epinephrine [6]) bind to receptors that couple to the G protein family members Gαi/o, leading to inhibition of adenylate cyclase and decreased intracellular cAMP levels. Following activation of GPCR signaling by agonist binding, signal termination depends on the GTPase activating protein (GAP) activity [7] of regulator of G protein signaling (RGS) proteins. Although the importance of GPCR to β-cell physiology and pharmacology is well documented [8], our understanding of the role of RGS proteins in the β cell is still incomplete.

Many of the 20 canonical mammalian RGS proteins are expressed in islet cell but only a few have been studied in the context of β-cell function. Both RGS2 and RGS4 negatively regulate insulin secretion in isolated mouse islets and in vivo [9; 10]. Using RNA silencing techniques in isolated mouse and human islets, we previously showed that RGS16 increases insulin secretion and β-cell proliferation [11]. Recent evidence suggests that the R7 RGS protein family member RGS9 may also have a role in the endocrine pancreas. Using transcriptomics and proteomics approaches, RGS9 was shown to be enriched in human islets [12]. Patch-seq experiments on human β cells identified *RGS9* as a gene positively correlated with total exocytosis, a proxy for hormone secretion [13]. *RGS9* knockdown decreased total exocytosis, thus providing evidence for involvement of RGS9 in insulin secretion. R7 RGS proteins contain a Dishevelled, Egl-10 and Pleckstrin (DEP) domain that is implicated in membrane targeting [14]. Like all R7 RGS protein family members, RGS9 forms an obligatory heterodimer with the atypical G protein Gβ5 via binding to its G protein γ-like (GGL) domain, that is necessary for protein stability [15]. Accordingly, *Gnb5* (Gβ5) knockout mouse islets exhibit defective glucose-stimulated insulin secretion (GSIS) [16; 17]. However, the precise role of individual R7 RGS proteins in this context is unknown.

This study aimed to determine whether RGS9 controls insulin secretion in rodent islets. As RGS mapping studies indicate that RGS9 acts exclusively on Gαi/o [18], we hypothesized that RGS9 enhances insulin secretion by alleviating inhibitory signaling in the β cell [3].

## 2. MATERIALS AND METHODS

### 2.1 Reagents and solutions

RPMI-1640 and FBS were obtained from Life Technologies Inc. (Burlington, ON, Canada). Penicillin/streptomycin was acquired from Multicell Wisent Inc (Saint-Jean-Baptiste, QC, Canada). FA-free BSA was obtained from Equitech-Bio (Kerrville, TX, USA). Insulin radioimmunoassay (RIA) kits were obtained from MilliporeSigma (Cat# RI-13K, RRID:AB_2884035). Adenoviruses expressing two distinct short hairpin (sh) RNAs against *Rgs9*, shRgs9(A) (Cat # shADV-270502) and shRgs9(B), and control scrambled shRNA (shScr), with GFP reporter were obtained from Vector Biolabs (Malvern, PA). All of the other reagents were acquired from MilliporeSigma unless otherwise specified.

### 2.2 Animals

Animals were handled in accordance with the Canadian Council on Animal Care guidelines for the care and use of laboratory animals. Mice were housed under controlled temperatures on a 12-h light/dark cycle with unrestricted access to water and standard laboratory chow. B6.129S1-*Rgs9^tm1Citb^*/Mmnc mice (*Rgs9^+/ΔDEP^*) on a mixed C57Bl/6JN background were purchased from the MMRRC (RRID:MMRRC_031829-UNC). Mice carrying the wild-type NNT allele were selected for breeding of experimental animals [19]. Experimental animals (*Rgs9^+/+^* and *Rgs9^ΔDEP/ΔDEP^*) were generated from crosses between *Rgs9^+/ΔDEP^*mice. Genotyping primers are listed in Supplementary Table 1. Animals were born at the expected Mendelian ratios. C57BL/6N male mice were purchased from Charles River (Saint-Constant, QC, Canada).

### 2.3 Fluorescence mRNA in situ hybridization

mRNAs were visualized by fluorescence in situ hybridization using an RNAscope Fluorescent Multiplex Kit (Advanced Cell Diagnostics, Hayward, CA) on fixed and frozen 12-week-old male C57Bl/6N pancreatic cryosections. Mouse pancreata were prepared as described in protocols.io (dx.doi.org/10.17504/protocols.io.e6nvw9367gmk/v1). Briefly, pancreata were fixed overnight in 4% paraformaldehyde and cryoprotected overnight in 30% sucrose. Tissues were then embedded in OCT, frozen, sectioned at 8 μm, and mounted on Superfrost Plus slides (Life Technologies, Carlsbad, CA). The following RNAscope probes were used: Mm-*Rgs9* (Cat # 468871), Mm-*Ins2*-O1-C2 (Cat # 497811-C2), Mm-*Sst*-C4 (Cat # 404631-C4), and Mm-*Gcg*-C3 (Cat # 400601-C3). Hybridization and fluorescent detection were performed according to the manufacturer’s instructions. 20 X and 63 X images were acquired with an inverted confocal microscope (Leica Microsystem, Mannheim, Germany).

### 2.4 Islet isolation and culture

Islets were isolated from 8-10 week-old male mice by collagenase digestion and dextran density gradient centrifugation as previously described [20] and allowed to recover overnight in RPMI 1640 supplemented with 10% (wt/vol) FBS, 100 U/ml of penicillin/streptomycin, and 11 mM of glucose (complete media).

### 2.5 Flow cytometric sorting of mouse α, β and δ cells

Flow cytometric sorting of mouse islet α, β and δ cells was performed as described [21]. Briefly, islets were dispersed in Accutase (Innovative Cell Technologies, Inc., San Diego, CA), incubated with antibodies and labeled with LIVE/DEAD Fixable Aqua vivid (405 nm) Dead Cell Stain (BD Biosciences, San Jose, CA). Antibodies and dilutions are indicated in Supplementary Table 2. Cell-sorting was performed using a FACSAria III (BD Biosciences). Sorted cell were collected directly in RLT buffer (Qiagen, Hilden, Germany) for real-time (q)PCR.

### 2.6 Adenoviral-infection and pseudo-islet generation

Batches of 400 islets were dispersed by gentle pipetting for 10 min at 37 °C in Accutase (500 μl, Innovative Cell Technologies, San Diego, CA). Dissociated islet cells were placed in 24-well Aggrewell 800 plates (STEMCELL Technologies, Vancouver, BC, Canada), infected with adenovirus at 80-100 multiplicity of infection, and subjected to centrifugal-force-aggregation. Islet cell suspensions were prepared in complete RPMI 1640 media to create 1000-1500 cell pseudo-islets. Adenoviruses were removed after 24 h and the pseudo-islets were transferred to culture dishes at 72 h. Pseudo-islets were cultured for an additional 4 days in complete RPMI 1640 media before experimentation.

### 2.7 Immunofluorescence microscopy

Pseudo-islets were fixed for 4 hours in 4% paraformaldehyde (PFA) and cryoprotected overnight in 30% sucrose. The tissues were embedded in OCT compound, frozen, sectioned at 8 μm, and mounted on Superfrost Plus slides (Thermo Fisher Scientific, Waltham, MA). Antigen retrieval was performed using sodium citrate buffer (pH=6). Sections were stained for Insulin, Glucagon and SST and nuclei were stained with Hoechst 33342 (Thermo Fisher Scientific). Primary and secondary antibodies and dilutions are listed in Supplementary Table 3. Slides were mounted using Vectashield mounting medium (Vector laboratories Inc, Newark CA) and images acquired using a Zeiss AX10 fluorescence microscope (Oberkochen, Germany).

### 2.8 RNA extraction and qPCR

Total RNA was extracted from intact islets or flow cytometry sorted α, β and δ cells using a RNeasy micro kit (Qiagen, Hilden, Germany). RNA was quantified by spectrophotometry using a NanoDrop 2000 (Life Technologies, RRID:SCR_018042) and 0.3 μg of RNA was reverse transcribed. qPCR was performed using a QuantiTect SYBR Green PCR kit (Qiagen). Data were normalized to cyclophilin A (*Ppia*) mRNA levels. qPCR primer sequences are listed in Supplementary Table 4.

### 2.9 Static incubations for insulin secretion

Islets and pseudo-islets were incubated in KRBH (pH 7.4) with 0.1% (w/v) FA-free BSA (KRBH/BSA) and 2.8 mM of glucose for 20 min. Then, triplicate batches of 20 islets/pseudo-islets each were distributed into each well of a 24-well plate and incubated for an additional 20 min in KRBH/BSA with 2.8 mM of glucose, followed by a 1-h static incubation in KRBH/BSA in the presence of 2.8 mM or 16.7 mM of glucose or 2.8 mM of glucose with 30 mM KCl. Secreted insulin was measured in the supernatant by RIA. Intracellular insulin content was measured after acid-alcohol extraction.

### 2.10 Statistical analyses

Data are expressed as mean ± SEM. Significance was tested using standard one-way ANOVA, Brown-Forsythe and Welch ANOVA and corrections, in cases of variance heterogeneity, two-way ANOVA with post hoc adjustment for multiple comparisons, and Paired and Unpaired Student’s t test, as appropriate, using GraphPad (RRID:SCR_000306). Tukey or Dunnett post hoc tests were performed as indicated in figure legends. p < 0.05 was considered significant.

### 2.11 Study approval

All procedures involving animals were approved by the Institutional Committee for the Protection of Animal at the Centre Hospitalier de l’Université de Montréal.

## 3. RESULTS

### 3.1 *Rgs9-1* is the major transcript isoform expressed in mouse pancreatic islets

Although *RGS9* expression has been reported to be enriched in human pancreatic islets, its cell type-specific localization within the rodent endocrine pancreas remains unclear. To address this, we performed RNAscope in situ hybridization on male mouse pancreatic sections using probes for *Rgs9* together with markers for β (Ins), α (Gcg), and δ cells (Sst). *Rgs9* transcripts were detected exclusively within the endocrine compartment and were absent from the exocrine pancreas (Fig. 1A). Qualitative analysis revealed a cell type-specific expression gradient, with the highest expression in δ cells, intermediate expression in β cells, and the lowest expression in α cells. This distribution is consistent with previously published islet transcriptomic datasets from the Huising laboratory [22].

**Figure 1.**
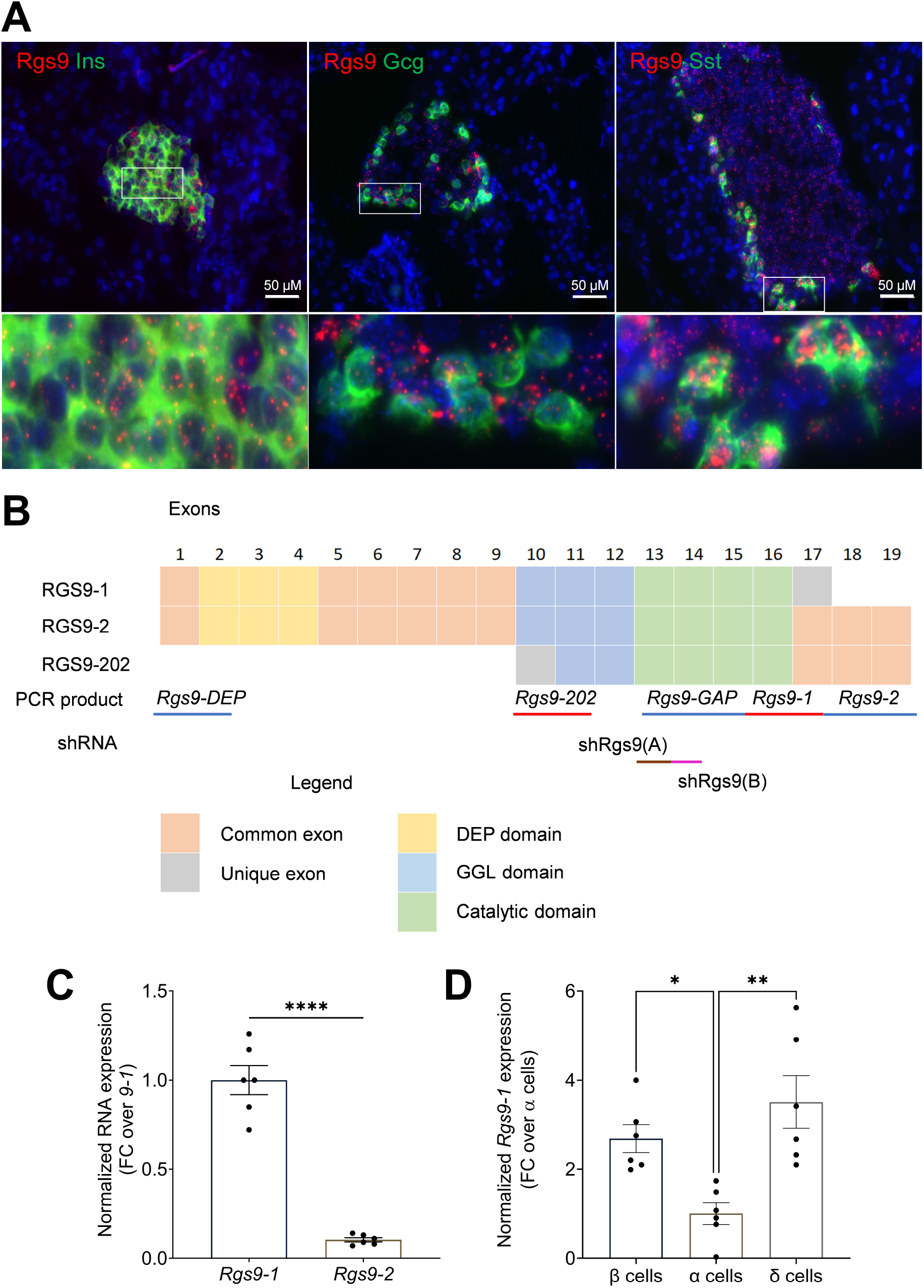
*Rgs9* transcript variants and endocrine cell-specific expression in mouse pancreatic islets. (A) Representative RNAscope in situ hybridization of mouse pancreatic sections showing *Rgs9* (red) together with *Ins2*, *Gcg*, or *Sst* (green). Nuclei were counterstained with DAPI (blue). (B) Schematic representation of the *Rgs9* transcript variants, domain structure, and the location of the qPCR amplicons and shRNA target sequences. (C) Relative mRNA expression of the *Rgs9-1* and *Rgs9-2* splice variants in isolated mouse islets. Transcripts were normalized to cyclophilin A and presented as the fold change (FC) over *Rgs9-1.* (D) Relative *Rgs9-1* mRNA expression in fluorescence-activated cell sorting (FACS)-purified β, α, and δ cells. Transcripts were normalized to cyclophilin A and presented as the fold change (FC) over α cells. Data are presented as mean ± SEM (N=6). Statistical significance was determined using an unpaired two-tailed Student’s t-test (C) or one-way ANOVA followed by Tukey’s multiple-comparisons test (D). *P < 0.05, **P < 0.01, ****P < 0.0001.

The *Rgs9* gene generates two major splice variants encoding the short RGS9-1 (484 amino acids) and long RGS9-2 (675 amino acids) isoforms [23] (Fig. 1B). Analysis of isoform-specific expression demonstrated that both transcripts are present in mouse islets, with *Rgs9-1* expressed at substantially higher levels than *Rgs9-2* (Fig. 1C). Given that RGS9-1 has been primarily described as a photoreceptor-specific isoform [24], its predominance in pancreatic islets was unexpected. To validate these findings, we quantified *Rgs9-1* expression in flow cytometry-sorted α, β, and δ cell populations. In line with the RNAscope data, *Rgs9-1* expression was higher in δ and β cells compared to α cells (Fig. 1D).

### 3.2 RGS9 supports insulin secretion

To investigate the role of RGS9 in insulin secretion, we infected mouse reaggregated pseudo-islets with adenoviruses expressing a shRNA targeting *Rgs9*. We used pseudo-islets rather than intact islets to enhance adenoviral transduction efficiency [25–27] (Fig. 2A). Control and adenovirally transduced pseudo-islets were of a comparable size and contained predominantly β and fewer δ and α cells, similar to intact islets (Fig. 2B). Pseudo-islets infected with two distinct shRgs9 constructs targeting GAP domain sequences (shRgs9(A) and shRgs9(B)) (Fig. 1B) exhibited a marked reduction in *Rgs9* mRNA expression compared with control (shScr) (Fig. 2C), confirming efficient gene silencing. We next assessed the impact of *Rgs9* depletion on insulin secretion. Both glucose- and KCl-induced insulin secretion were significantly reduced following knockdown with shRgs9(A), whereas shRgs9(B) produced a similar but non-significant reduction compared to controls (Figs. 2D&E). Total insulin content was comparable between shRgs9(A), shRgs9(B) and control infected pseudo-islets (Fig. 2F), indicating that *Rgs9* depletion impairs insulin secretion without significantly affecting insulin stores. Given its more robust effects, shRgs9(A) was used in subsequent experiments.

**Figure 2.**
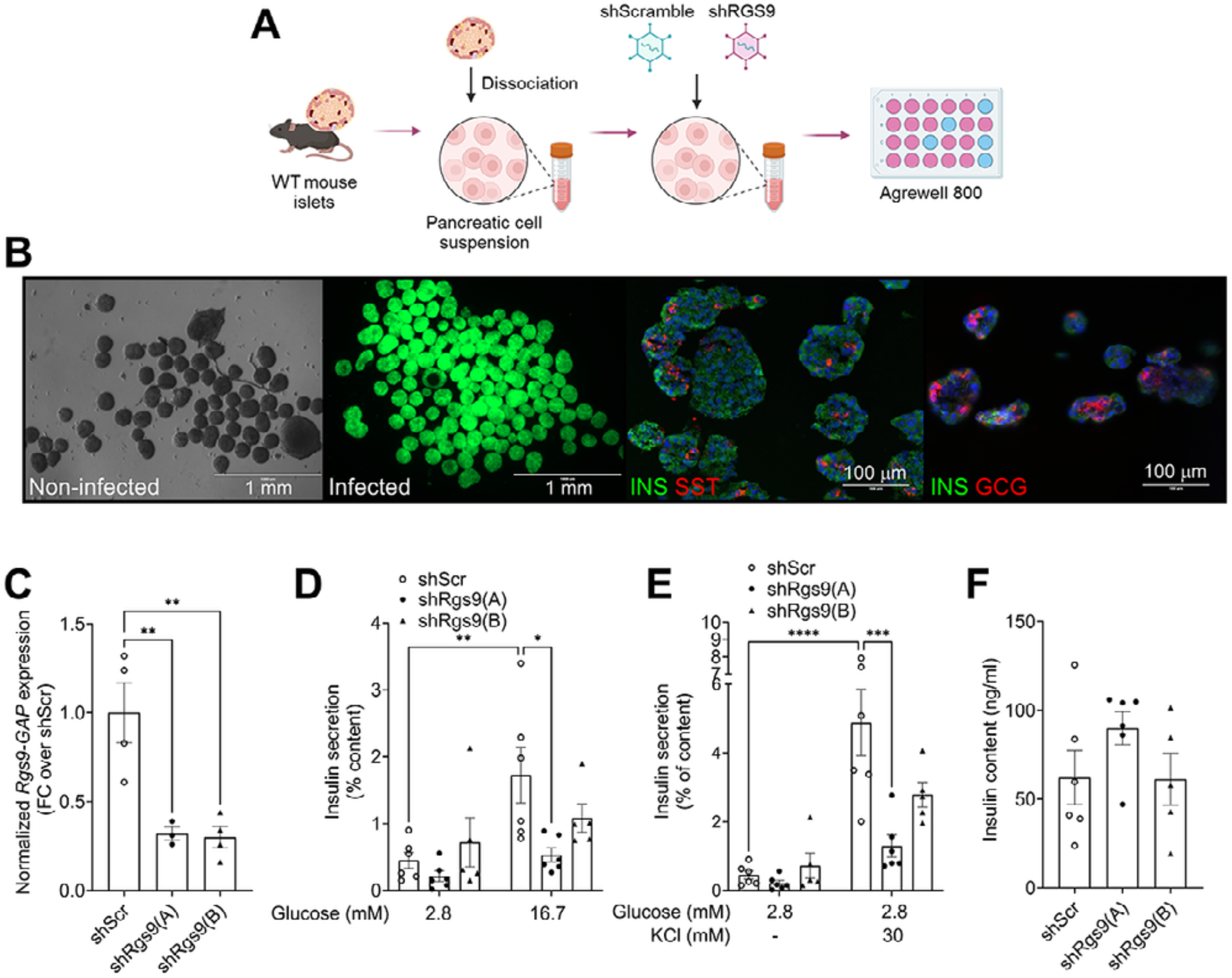
Acute *Rgs9* knockdown in mouse pseudo-islets impairs glucose- and depolarization-induced insulin secretion. (A) Schematic illustrating the generation of mouse pseudo-islets. Isolated islets were dissociated, infected with adenoviruses expressing shRNA, and reaggregated in AggreWell 800 plates. (B) (Left panels) Representative images of non-infected pseudo-islets and pseudo-islets infected with control (shSrc) adenovirus. GFP fluorescence reveals transduced cells. (Right panels) Pseudo-islets sections were immunostained for insulin (INS), glucagon (GCG) and somatostatin (SST), and DAPI. Scale bars, 1 mm and 100 μm. (C) Relative *Rgs9* mRNA expression, based on detection of *GAP* domain sequences (Fig. 1B), in pseudo-islets infected with shRgs9(A), shRgs9(B) or control (shScr) adenovirus. Transcripts were normalized to cyclophilin A and presented as the fold change (FC) over shScr. (D&E) Insulin secretion, expressed as a percentage of total insulin content, from pseudo-islets infected with shRgs9(A), shRgs9(B) or control (shScr) adenovirus following exposure to 2.8 or 16.7 mM glucose (D), and 2.8 mM glucose with or without 30 mM KCl (E). (F) Total insulin content (ng/ml) of pseudo-islets infected with shRgs9(A), shRgs9(B) or control (shScr) adenovirus. Data are presented as mean ± SEM (N=3-6). Statistical significance was determined by two-way ANOVA followed by Tukey’s multiple-comparisons test. *P < 0.05, **P < 0.01, ***P < 0.001, ****P < 0.0001.

### 3.3 Deletion of the RGS9 DEP domain in mice does not impair insulin secretion

To further investigate the role of RGS9 in insulin secretion, we used a *Rgs9* mutant mouse model in which exons 2–4, encoding the DEP domain (Fig. 1B), are deleted (*Rgs9^ΔDEP^*, [28]). Glucose- and KCl-stimulated insulin secretion were assessed in isolated islets from wild-type (*Rgs9^+/+^*) and mutant (*Rgs9^ΔDEP/ΔDEP^*) mice. Surprisingly, and in contrast to the results shown in Figs. 2D&E, no differences in insulin secretion were observed between control and mutant islets under either basal or stimulatory conditions (Fig. 3A), suggesting that deletion of the DEP domain does not alter β-cell function.

**Figure 3.**
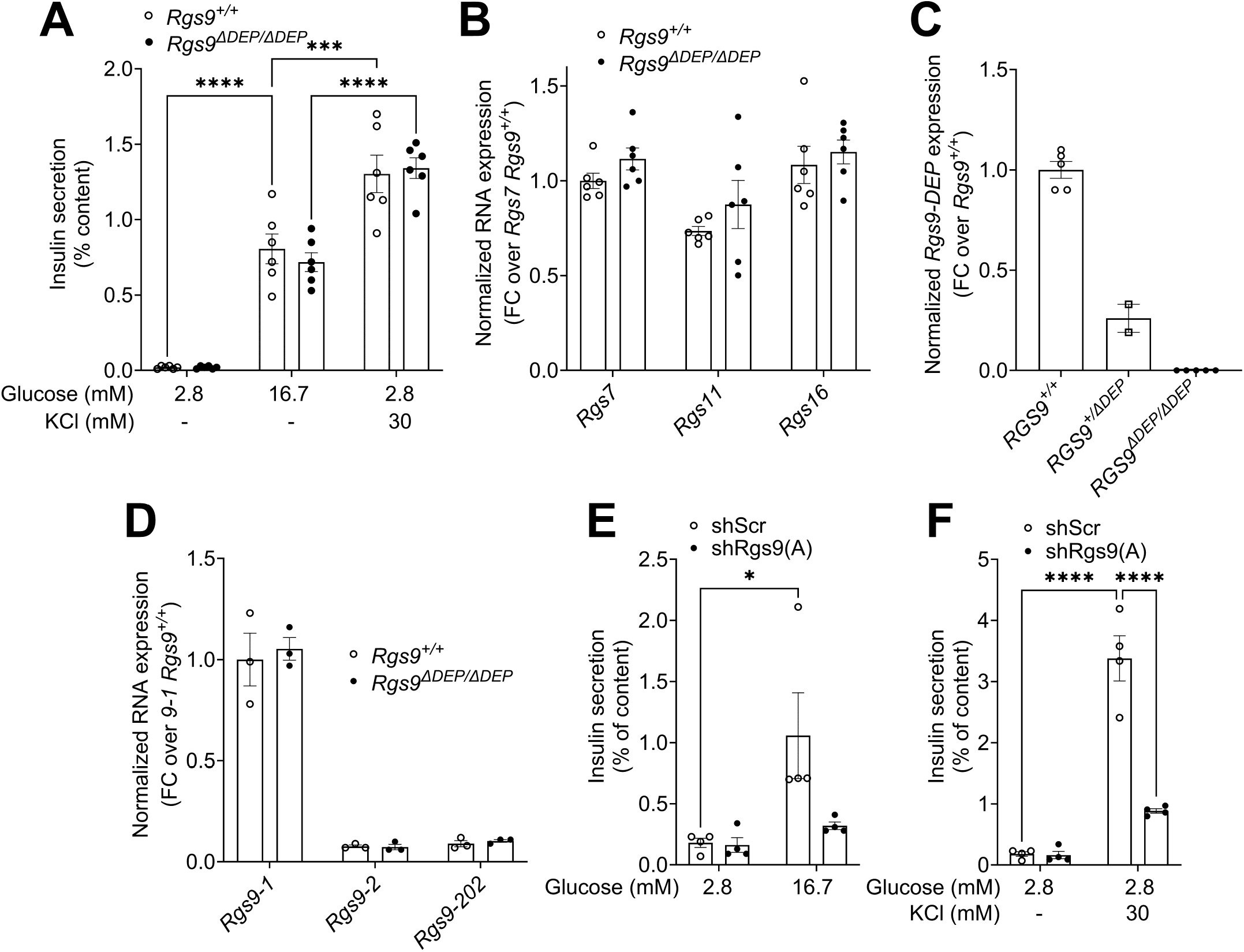
*Rgs9^ΔDEP/ΔDEP^* islets express functional *Rgs9* transcripts to maintain stimulated insulin secretion. (A) Insulin secretion, expressed as a percentage of total insulin content, from *Rgs9^+/+^*and *Rgs9^ΔDEP/ΔDEP^* islets following exposure to 2.8 mM glucose, 16.7 mM glucose, or 30 mM KCl. (B) Relative mRNA expression of *Rgs7*, *Rgs11*, and *Rgs16* in *Rgs9^+/+^* and *Rgs9^ΔDEP/ΔDEP^* islets. Transcripts were normalized to cyclophilin A and presented as the fold change (FC) over *Rgs7* in *Rgs9^+/+^.* (C) Relative mRNA expression of *Rgs9* transcripts carrying the *DEP* domain sequences in *Rgs9^+/+^*, *Rgs9^+/ΔDEP^* and *Rgs9^ΔDEP/ΔDEP^*islets. Transcripts were normalized to cyclophilin A and presented as the fold change (FC) over *Rgs9^+/+^.* (D) Relative mRNA expression of the *Rgs9-1*, *Rgs9-2*, and *Rgs9-202* transcript variants in *Rgs9^+/+^* and *Rgs9^ΔDEP/ΔDEP^* islets. Transcripts were normalized to cyclophilin A and presented as the fold change (FC) over *Rgs9-1* in *Rgs9^+/+^.* (E&F) Insulin secretion, expressed as a percentage of total insulin content, from *Rgs9^ΔDEP/ΔDEP^* pseudo-islets infected with shRgs9(A) or control (shScr) adenovirus following exposure to 2.8 or 16.7 mM glucose (E), and 2.8 mM glucose with or without 30 mM KCl (F). Data are presented as mean ± SEM (N=2-6). Statistical significance was determined by two-way ANOVA followed by Tukey’s multiple-comparisons test. *P < 0.05, ***P < 0.001, ****P < 0.0001.

The discrepancy between these findings and the reduced GSIS observed upon acute knockdown of RGS9 could be due to compensatory changes in the expression of other RGS proteins in the constitutive knockout model. To assess this possibility, we measured the expression of other members of the R7 family, *Rgs7* and *Rgs11*, together with *Rgs16*, which we previously showed to play a role in insulin secretion [11]. No significant differences in the expression of either *Rgs7*, *Rgs11*, or *Rgs16* was detected between wild-type and mutant islets (Fig. 3B), suggesting that these *Rgs* gene transcripts do not compensate for *Rgs9* deletion.

We next characterized *Rgs9* transcript expression in mutant islets. As expected, PCR analysis confirmed the absence of transcripts containing exons 2–4, corresponding to the deleted DEP domain (Fig. 1B), in *Rgs9^ΔDEP/ΔDEP^* islets compared with *Rgs9^+/+^* and *Rgs9^+/ΔDEP^* controls (Fig. 3C). However, despite the absence of the DEP domain, expression of transcripts of *Rgs9-1*, *Rgs9-2* and *Rgs9-202* (Fig. 1B), a shorter *Rgs9* transcript isoform predicted to retain an intact catalytic domain, were comparable between wild-type and mutant islets (Fig. 3D).

To confirm that complete deletion of RGS9 transcripts inhibits insulin secretion, dissociated *Rgs9^ΔDEP/ΔDEP^* islets were infected with either shRgs9(A) or a control adenovirus and reaggregated into pseudo-islets. RGS9 knockdown in *Rgs9^ΔDEP^*islets significantly reduced insulin secretion in response to both high glucose (16.7 mM) and KCl (30 mM) compared with controls (Fig. 3E&F), reproducing the phenotype observed in wild-type pseudo-islets (Fig. 2D&E). Collectively, these findings indicate 1-that the *Rgs9^ΔDEP^* mutant mouse does not represent a complete loss-of-function model in islets, and 2-that the DEP domain is dispensable for RGS9 regulation of insulin secretion.

## 4. DISCUSSION

In the present study, we identify RGS9 as a previously unrecognized regulator of pancreatic islet function. Our results show that 1-*Rgs9* is expressed in the endocrine pancreas and is enriched in β and δ cells with minimal expression in α cells; 2-acute knockdown of *Rgs9* impairs both glucose- and KCl-stimulated insulin secretion without affecting insulin content; and 3-deletion of the DEP domain in *Rgs9^ΔDEP/ΔDEP^* mouse islets fails to reproduce this phenotype. We conclude that RGS9 promotes insulin secretion in β cells that the DEP domain is dispensible for this effect.

Previous single-cell transcriptomic analyses identified *Rgs9* as an islet-enriched gene and showed preferential expression in β and δ cells [22; 29; 30]. Using RNAscope and fluorescence-sorted endocrine cell populations, we demonstrate that *Rgs9* expression is highest in mouse β and δ cells. We also found that *Rgs9-1* is the predominant splice variant expressed in mouse islets. This observation is surprising because *Rgs9-1* has long been considered a retina-specific isoform, whereas *Rgs9-2* predominates in the brain [31; 32]. These findings suggest that *Rgs9* may have previously unrecognized functions outside the nervous system and highlight the importance of considering isoform-specific regulation in endocrine tissues. Unfortunately, we were unable to confirm protein expression because commercially available RGS9 antibodies lacked sufficient specificity and sensitivity.

A striking finding of this study is the discrepancy between the acute knockdown of *Rgs9* and a genetic mutant model. Although acute depletion of *Rgs9* via shRNA-mediated knockdown consistently impaired glucose- and KCl-stimulated insulin secretion, islets isolated from *Rgs9^ΔDEP/ΔDEP^* mice displayed unaltered secretory responses. Phenotypic differences between acute knockdown and genetic mutants are increasingly recognized and may arise from transcriptional adaptation whereby related genes are upregulated to compensate for the targeted gene deletion [33; 34]. Here, we found no evidence for compensatory changes in the expression of other R7 family *Rgs* genes including *Rgs7* and *Rgs11* or the β-cell-enriched *Rgs16,* which we showed previously to promote insulin secretion [11]. However, we cannot exclude compensation by other RGS family proteins or by proteins with overlapping GAP activity toward Gαi/o, at either the transcriptional or post-transcriptional level. Furthermore, we were unable to evaluate the expression of Gβ5, an obligate binding partner required for the stability and activity of R7 family proteins [35]. Nevertheless, the discrepancy with acute *Rgs9* knockdown is likely explained by the persistence of transcripts encoding the catalytic RGS domain despite deletion of exons 2–4 encompassing the DEP domain in *Rgs9^ΔDEP/ΔDEP^*mice. While we cannot conclude that these transcripts produce a functional protein, their persistence indicates that this model is not a complete null allele in pancreatic islets. In line with this possibility, shRNA-mediated knockdown of *Rgs9^ΔDEP^* transcripts in islets impaired insulin secretion, suggesting that the remaining transcripts are functional. Of note, these transcripts lack the DEP domain, suggesting that DEP domain-deficient RGS9 is functional in islets. Although the RGS9 DEP domain is necessary for membrane localization via membrane-anchoring proteins like R7BP and R9AP [36–38], in the absence of the DEP domain, the GAP domain remains intact and functional [39].

We demonstrated that shRNA-mediated knockdown of *Rgs9* impairs glucose- and KCl-stimulated insulin secretion without affecting insulin content. These data are consistent with previous findings in human β cells demonstrating that *RGS9* expression positively correlates with exocytosis and that siRNA-mediated knockdown diminishes glucose-stimulated exocytosis [13]. Furthermore, deletion of *Gnb5*, the gene encoding the atypical G protein subunit Gβ5, in islets leads to the degradation of all R7 RGS proteins and a decrease in GSIS [17]. As RGS9 would also be expected to be degraded in the absence of Gβ5, the phenotype of *Gnb5^-/-^*may in part be due to the loss of RGS9. Given that RGS9 acts exclusively on Gαi/o [18], it is likely that RGS9 promotes insulin secretion by mitigating inhibitory signaling in β cells, although this hypothesis remains to be directly tested. Pancreatic β cells express several inhibitory GPCR including the SST, α2A-adrenergic, galanin, and dopamine receptors capable of suppressing cAMP production and insulin exocytosis [40; 41]. Because RGS9 accelerates GTP hydrolysis on Gαi/o, loss of RGS9 would be expected to prolong signaling downstream of these receptors. Whether RGS9 selectively regulates individual GPCR pathways or instead acts as a broader modulator of inhibitory Gαi/o signaling remains to be determined.

In conclusion, our study identifies RGS9 as a novel regulator of pancreatic islet function where it contributes to the regulation of insulin secretion and intra-islet communication. Given that RGS9 is an important modulator of Gαi/o activity with a potential links to numerous inhibitory GPCR in the β cell, it will be important to investigate whether *Rgs9* dysregulation contributes to islet dysfunction in obesity and type 2 diabetes.

## ACKNOWLEDGEMENTS

This study was supported by a Discovery Grant from the Natural Sciences and Engineering Research Council of Canada (RGPIN-2016-03952 to V.P.) and the U.S. National Institutes of Health (R01-DK-132597 to V.P.). L.R. was supported by a fellowship from the Société Francophone du Diabète and a travel award from the Quebec Cardiometabolic Health, Diabetes and Obesity Research Network. We thank Sabrina Sedano-Benitez and Gaël Dulude from the CRCHUM for valuable technical assistance, and Raphaël Scharfmann and Alexis Fouque (INSERM, Paris, France) for sharing unpublished cell-sorting protocols.

## 5. AUTHOR CONTRIBUTIONS

**Laura Reininger**: Conceptualization, methodology, investigation, formal analysis, and writing original draft. **Sarah Ferragne**: Methodology and investigation. **Scott Campbell**: Methodology and investigation. **Caroline Tremblay**: Methodology and investigation. **Mélanie Ethier**: Methodology. **Julien Ghislain**: Conceptualization, validation, writing, review, editing, supervision, and project administration. **Vincent Poitout**: Conceptualization, validation, writing, review, editing, supervision, funding acquisition, and project administration.

## Notes

### Competing Interest Statement

The authors have declared no competing interest.

