## Supplemental tables for "The regulator of G protein signaling RGS9 supports insulin secretion in male mouse islets"

**SUPPLEMENTARY TABLES FOR REININGER ET AL.**

**Supplementary Table 1: Primers used for genotyping.**

| **Primer** | **Sequence** |
| --- | --- |
| RGS9R | CACTGAGGCATGACCCCCAC |
| RGS9F | CCTGCCTCTCCTGCCTTTCG |
| Neo3 | CGGCGAGGATCTCGTCGTGACCCA |
| NeoT | GAACAAACTACCCAACACCCGTGCGT |

**Supplementary Table 2: Sources of antibodies used for flow cytometry.**

| **Fluorophore** | **Antibody** | **Dilution** | **Reference** |
| --- | --- | --- | --- |
| PerCP/Cyanine5.5 | Anti-mouse TER-119 | 1/100 | BioLegend; 116227, [RRID:AB_893638](https://rrid.site/data/record/nif-0000-07730-1/AB_893638/resolver?q=AB_893638&l=AB_893638&i=rrid:ab_893638-2183982) |
| PerCP/Cyanine5.5 | Anti-mouse CD31 | 1/100 | BioLegend; 102522, [RRID:AB_2566761](https://rrid.site/data/record/nif-0000-07730-1/AB_2566761/resolver?q=AB_2566761&l=AB_2566761&i=rrid:ab_2566761-2502217) |
| PerCP/Cyanine5.5 | Anti-mouse CD45 | 1/100 | BioLegend; 103131, [RRID:AB_893344](https://rrid.site/data/record/nif-0000-07730-1/AB_893344/resolver?q=AB_893344&l=AB_893344&i=rrid:ab_893344-2183739) |
| APC/Fire™ 750 | Anti-mouse CD24 | 1/100 | BioLegend; 101840, [RRID:AB_2650876](https://rrid.site/data/record/nif-0000-07730-1/AB_2650876/resolver?q=AB_2650876&l=AB_2650876&i=rrid:ab_2650876-2586332) |
| BV605 | Anti-mouse CD326 (Ep-CAM) | 1/100 | Biolegend; 118227, [RRID:AB_2563984](https://rrid.site/data/record/nif-0000-07730-1/AB_2563984/resolver?q=AB_2563984&l=AB_2563984&i=rrid:ab_2563984-2499440) |
| PE-Cyanine7 | Anti-mouse CD71 | 1/100 | BioLegend; 113811, [RRID:AB_2203383](https://rrid.site/data/record/nif-0000-07730-1/AB_2203383/resolver?q=AB_2203383&l=AB_2203383&i=rrid:ab_2203383-1656939) |
| PE | Anti-human/mouse CD49f | 1/200 | BioLegend; 313611, [RRID:AB_893374](https://rrid.site/data/record/nif-0000-07730-1/AB_893374/resolver?q=AB_893374&l=AB_893374&i=rrid:ab_893374-2183768) |
| BV421 | Anti-mouse/rat CD81 | 1/100 | BD biosciences; 740060, [RRID:AB_2739825](https://rrid.site/data/record/nif-0000-07730-1/AB_2739825/resolver?q=AB_2739825&l=AB_2739825&i=rrid:ab_2739825-2675281) |

**Supplementary Table 3: Sources of antibodies used for immunohistochemistry.**

| **Antibody** | **Dilution** | **Reference** |
| --- | --- | --- |
| Anti-human Insulin | 1/500 | Agilent; A0564  [RRID:AB_10013624](https://rrid.site/data/record/nif-0000-07730-1/AB_10013624/resolver?q=DAKO%20A0564&l=DAKO%20A0564&i=rrid:ab_10013624-2552625) |
| Anti-mouse Somatostain | 1/200 | Abcam; ab108456, [RRID:AB_11158517](https://rrid.site/data/record/nif-0000-07730-1/AB_11158517/resolver?q=AB_11158517&l=AB_11158517&i=rrid:ab_11158517-1221777) |
| Anti-pig Glucagon | 1/400 | Abcam; ab10988,  [RRID:AB_297642](https://rrid.site/data/record/nif-0000-07730-1/AB_297642/resolver?q=AB10988&l=AB10988&i=rrid:ab_297642-81344) |
| Cy3 Goat anti-mouse | 1/400 | Thermo Fisher Scientific; A10521, [RRID:AB_10373848](https://rrid.site/data/record/nif-0000-07730-1/AB_10373848/resolver?q=AB_10373848&l=AB_10373848&i=rrid:ab_10373848-2469486) |
| Alexa Fluor 488 Donkey anti-rabbit | 1/500 | Thermo Fisher Scientific; A21206, [RRID:AB_2535792](https://rrid.site/data/record/nif-0000-07730-1/AB_2535792/resolver?q=AB_2535792&l=AB_2535792&i=rrid:ab_2535792-2471248) |
| Alexa Fluor 488 Donkey Anti-Guinea Pig | 1/300 | Jackson ImmunoResearch Labs; 706-545-148  [RRID:AB_2340472](https://rrid.site/data/record/nif-0000-07730-1/AB_2340472/resolver?q=706-545-148&l=706-545-148&i=rrid:ab_2340472-2276554) |

**Supplementary Table 4: Primers used for qPCR.**

| **RNA** | **Forward** | **Reverse** |
| --- | --- | --- |
| *Rgs9-DEP* | CTTGGGGGTTTCCCCTTTCG | TCTGCATGTCCTTCACCAGG |
| *Rgs9-GAP* | GATGCGAGTGGAGAGATGGG | GGCCAGGAACAGCTTGTAGA |
| *Rgs9-1* | GGAAACCACCAAGAGAAGCTCCA | TGAGTTGGCTCCTGCGGTC |
| *Rgs9-2* | GTTTTTGCCAGGCTCTCACC | GGTCATCCGAGTCCATCAGG |
| *Rgs9-202* | ACCGACTTAATGGCACCGAT | GACGAGAACTGCTCGCTGTA |
| *Ppia* | TCTGAGCACTGGAGAGAAAGG | TTCTCTCCGTAGATGGACCTG |
